# Genome-wide association analyses of sleep disturbance traits identify new loci and highlight shared genetics with neuropsychiatric and metabolic traits

**DOI:** 10.1101/082792

**Authors:** Jacqueline M. Lane, Jingjing Liang, Irma Vlasac, Simon G. Anderson, David A. Bechtold, Jack Bowden, Richard Emsley, Shubhroz Gill, Max A. Little, AnneMarie I. Luik, Andrew Loudon, Frank A.J.L. Scheer, Shaun M. Purcell, Simon D. Kyle, Deborah A. Lawlor, Xiaofeng Zhu, Susan Redline, David W. Ray, Martin K. Rutter, Richa Saxena

**Author notes:** Equal Contribution (Martin K. Rutter, Richa Saxena). Corresponding author, R. Saxena, Center for Human Genetic Research, Massachusetts General Hospital, 185 Cambridge Street, CPZN 5.806, Boston, MA, 02114, USA.

## Abstract

Chronic sleep disturbances, associated with cardio-metabolic diseases, psychiatric disorders and all-cause mortality^1,2^, affect 25–30% of adults worldwide^3^. While environmental factors contribute importantly to self-reported habitual sleep duration and disruption, these traits are heritable^4–9^, and gene identification should improve our understanding of sleep function, mechanisms linking sleep to disease, and development of novel therapies. We report single and multi-trait genome-wide association analyses (GWAS) of self-reported sleep duration, insomnia symptoms including difficulty initiating and/or maintaining sleep, and excessive daytime sleepiness in the UK Biobank (n=112,586), with discovery of loci for insomnia symptoms (near *MEIS1, TMEM132E, CYCL1, TGFBI* in females and *WDR27* in males), excessive daytime sleepiness (near *AR/OPHN1*) and a composite sleep trait (near *INADL* and *HCRTR2*), as well as replication of a locus for sleep duration (at *PAX-8*). Genetic correlation was observed between longer sleep duration and schizophrenia (r_G_=0.29, *p*=1.90x10^−13^) and between increased excessive daytime sleepiness and increased adiposity traits (BMI r_G_=0.20, *p=*3.12x10^−09^; waist circumference r_G_=0.20, *p=*2.12x10^−07^).

---

Rather than being ‘secondary’, evidence suggests disordered sleep may play an important role in the etiology and maintenance of physical and mental health^1,2^. Heritability has been estimated at ~40% for sleep duration^4,6–8^, 25–45% for insomnia^9^ and 17% for excessive daytime sleepiness^9^, but few genetic factors are known. A Mendelian short sleep mutation in *BHLHE41* (P385R) has been identified, and confirmed in mouse models^10^. GWAS for sleep duration have been reported^11–14^, but only an association at the *PAX8* locus reached genome-wide significance and was confirmed across ethnic groups^12^. There are several reported loci for restless legs syndrome (RLS) and narcolepsy, but no known robust genetic loci for insomnia symptoms or excessive daytime sleepiness^15,16^.

We and others performed a GWAS for chronotype in the UK Biobank^17,18^ and a 23&me participant sample^19^. To identify genetic variants that contribute to self-reported sleep duration, insomnia symptoms, and excessive daytime sleepiness and link them with other conditions, we performed GWAS using phenotype measures in UK Biobank participants of European ancestry. Variation in sleep duration, insomnia symptoms and excessive daytime sleepiness was associated significantly with age, sex, principal components of ancestry (PCs), genotyping array, depression, psychiatric medication use, self-reported sleep apnea, and BMI (**Supplementary Table 1)**, as previously reported^20–23^. Together age, sex, and PCs explained 0.4%, 3.0% and 1.3% of variation in sleep duration, insomnia symptoms, and excessive daytime sleepiness respectively. Strong and significant pair-wise phenotypic correlation was seen between the traits overall and within each sex, with limited correlation observed with chronotype. (**Fig. 1a; Supplementary Fig. 1**).

**Figure 1.**
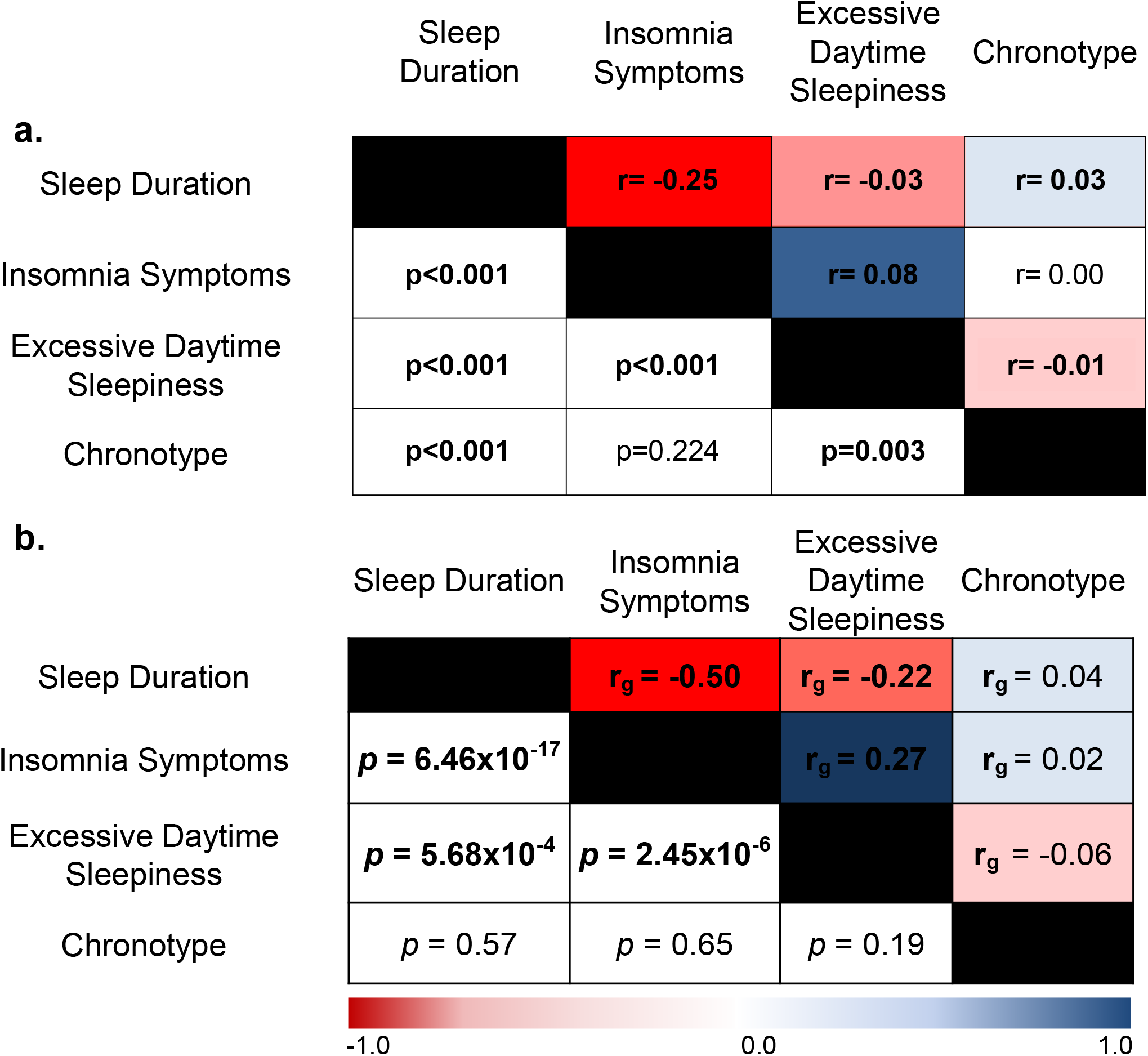
Sleep traits are phenotypically and genetically correlated. **a.** Phenotypic correlation between the reported sleep traits, using Spearman correlation (r). **b.** Genetic correlation (r_G_) between the reported sleep traits, using LD-score regression^67^. Color scale represents the strength of the correlation. Chronotype ranges from extreme morning types to extreme evening types.

GWAS analyses of sleep duration, insomnia symptoms and excessive daytime sleepiness were performed using linear/logistic regression adjusting for age, sex, 10 PCs and genotyping array. Nine genome-wide significant (*p*<5x10^−8^) and 14 suggestive (*p*<5x10^−7^ to *p*=5x10^−8^) loci were identified (**Fig. 2, Table 1, Supplementary Figs. 2 and 3**). For sleep duration (n=111,975), the strongest association was observed at the *PAX-8* locus (rs62158211T, β(se)=2.34(0.30) mins/allele, *p*=4.7x10^−14^, effect allele frequency (EAF) 0.213, **Fig. 2a**), confirming a previously reported association (r^2^=0.96, D’=1 to lead SNP rs1823125 in 1KG CEU)^12^. For insomnia symptoms (n=32,155 cases, 26,973 controls), significant associations were observed within *MEIS1* (rs113851554T, OR [95%CI]=1.26[1.20-1.33], *p*=9.1x10^−19^, EAF 0.057, **Fig. 2b**), a homeobox gene implicated in motor neuron connectivity in Drosophila^24,25^, retinal and lens development in mouse^26^, and Substance P expression in the amygdala^27^, near *TMEM132E* (rs145258459C, 1.23[1.13–1.35], *p*=2.1x10^−8^, EAF 0.983, **Fig. 2c**), a gene family with roles in brain development^28^, panic/anxiety^29^ and bipolar disorder^30^, suggesting a link between insomnia symptoms and an underlying broader sensitivity to anxiety and stress, and near *CYCL1* (rs5922858G, OR [95%CI]=1.12[1.07-1.16], *p=*1.28 x10^−8^, EAF 0.849, **Fig 2d**) a locus previously associated (*p*=10^−6^) with alcohol dependence co-morbid with depressive symptoms^31^. Sex-stratified analyses identified an additional female-specific signal near *TGFBI (*rs3792900C 1.10[1.07-1.14], *p*=2.16x10^−8^, EAF 0.470; **Table 1, Supplementary Fig. 3q, 3r, Supplementary Table 2**), an extracellular matrix protein responsible for human corneal dystrophy^32^ and a male-specific signal near *WDR27,* a scaffold protein (rs13192566G OR [95%CI]=1.14[1.09-1.20], *p*=3.2x10^−8^, EAF 0.860**)(Table 1**, **Supplementary Fig. 3s, 3t, 4**; **Supplementary Table 2**). Independent associations at both loci are observed with type 1 diabetes, suggesting an immune role^33–35^. For excessive daytime sleepiness (n=111,648), we identified a signal near the androgen receptor *AR* (rs73536079T, β=0.634, *p=*3.94x10^−8^, EAF 0.002, **Fig. 3e)**, with no sex-specific effects. Secondary analyses after additional adjustment for depression or BMI identified a signal near *ROBO1,* (depression adjustment n=107,440, rs182765975T, beta=0.099, p=3.33x10^−8^, EAF 0.003, **Table 1, Supplementary Figure 3o**), a neuronal axon guidance receptor previously implicated in dyslexia^36^, and a signal near another member of the TMEM132 family, *TMEM132B (*BMI adjustment n=75,480, rs142261172A, β=0.106, *p=*9.06x10^−9^, EAF 0.004, **Table 1, Supplementary Figure 3p**). Conditional analyses did not identify independent association signals (**Supplementary Table 3**). Sensitivity analyses adjusting for factors influencing sleep traits, including self-reported sleep apnea, depression, psychiatric medication use, smoking, socio-economic status, employment status, marital status, and snoring did not significantly alter results for primary association signals (**Supplementary Table 4**).

**Table 1.**
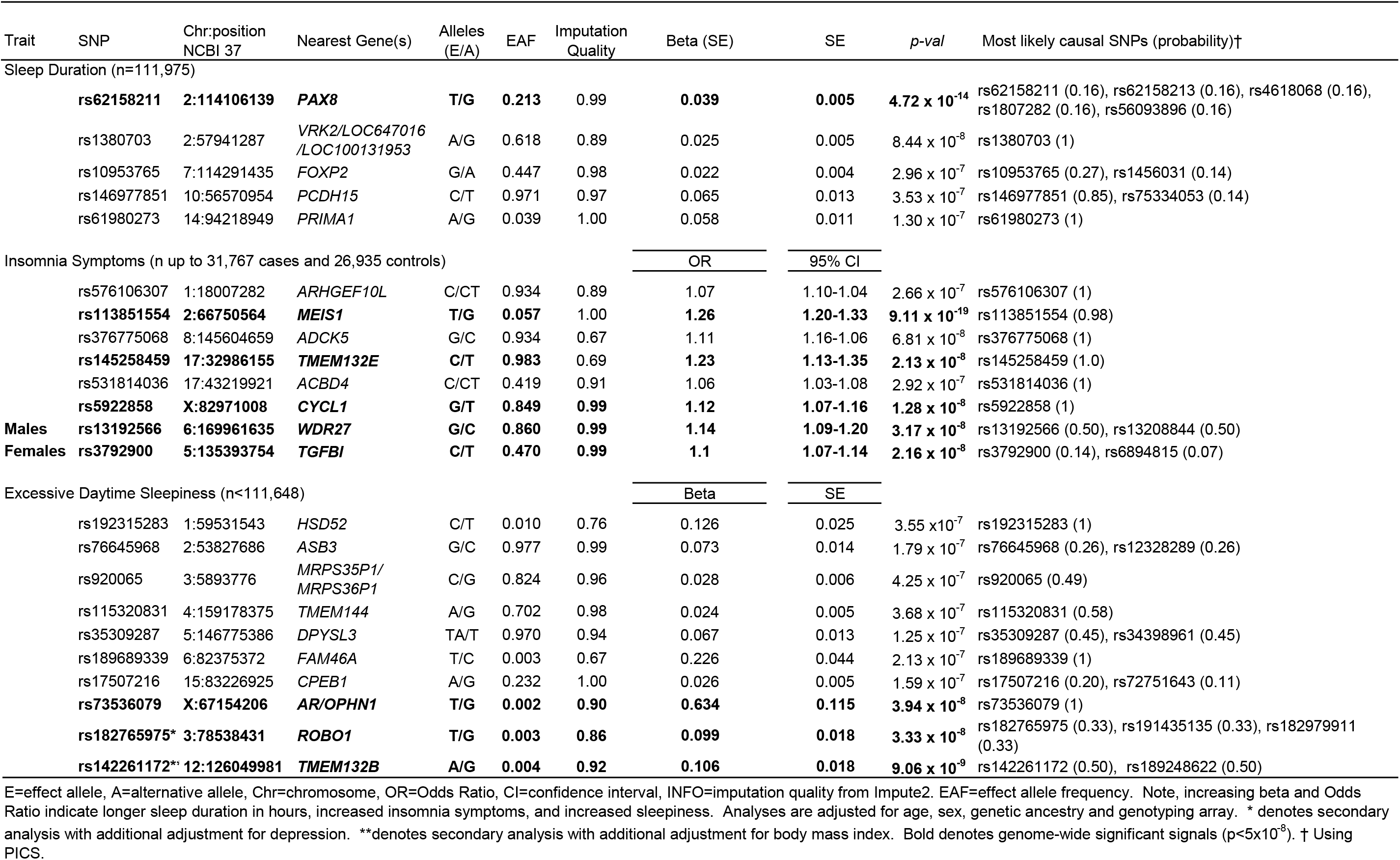
Genome-wide significant (p<5x10^−8^) and suggestive (p<5x10^−7^) loci associated with sleep duration, insomnia symptoms, and excessive daytime sleepiness in subjects of European ancestry in the UKBiobank.

**Figure 2.**
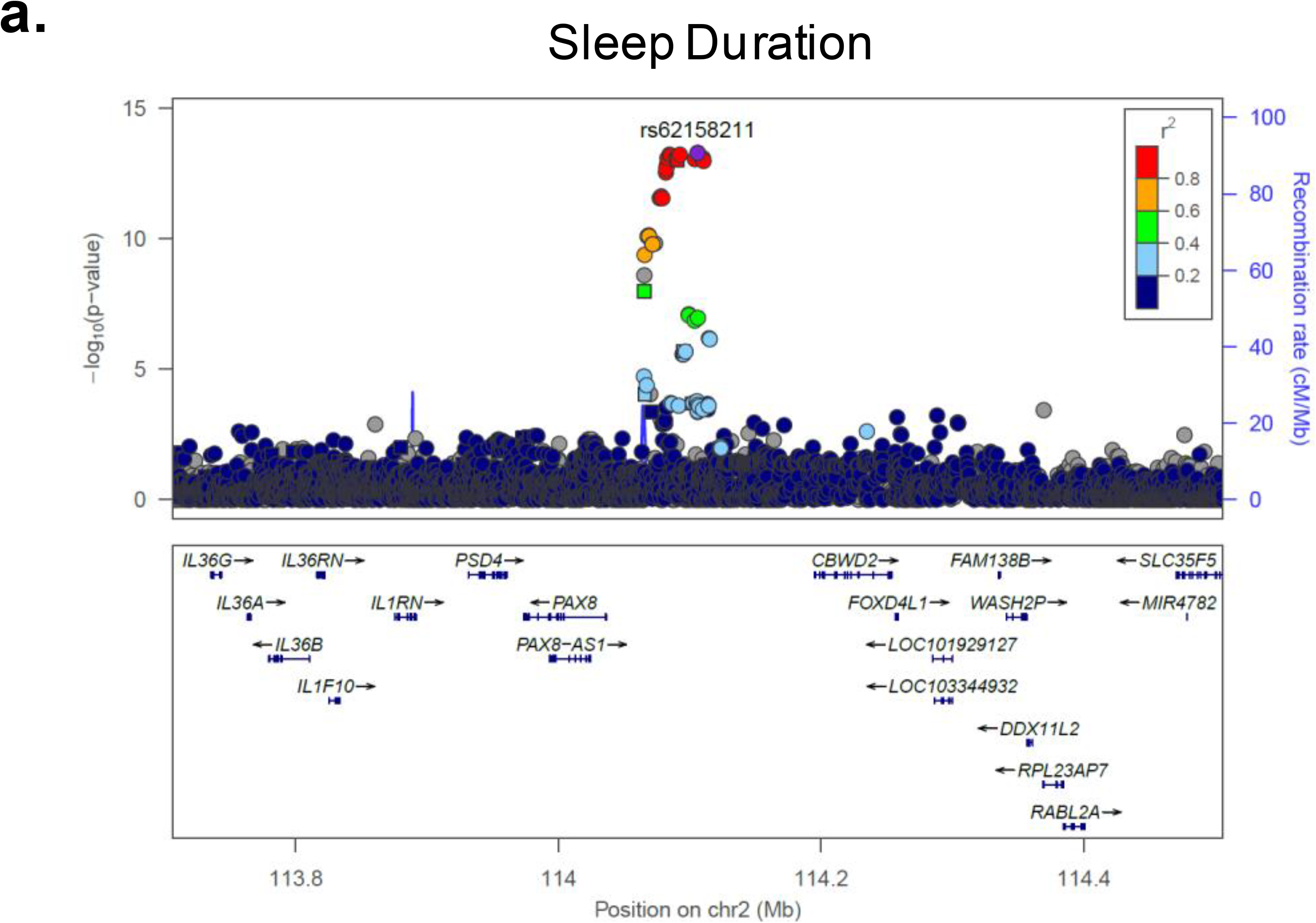
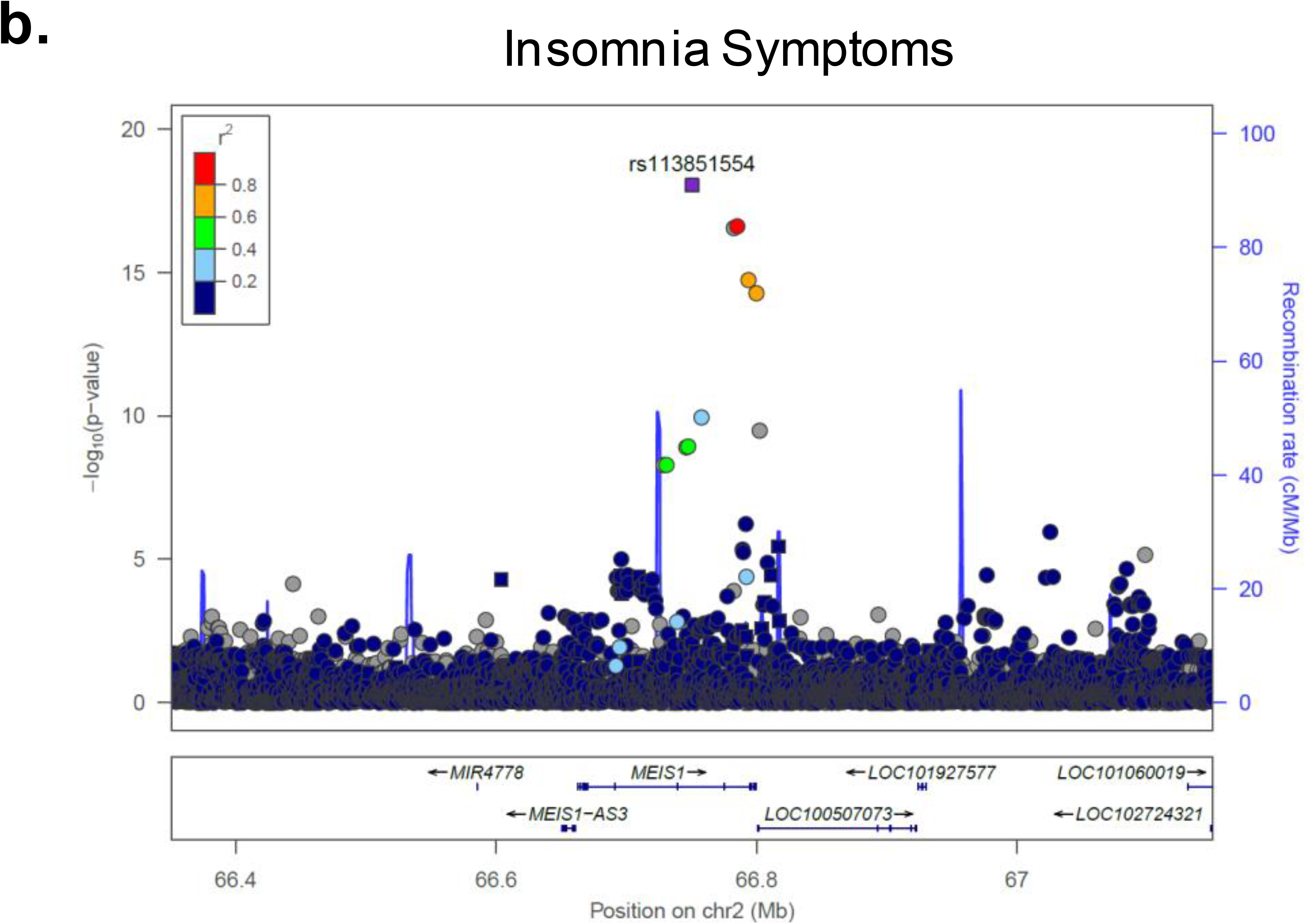
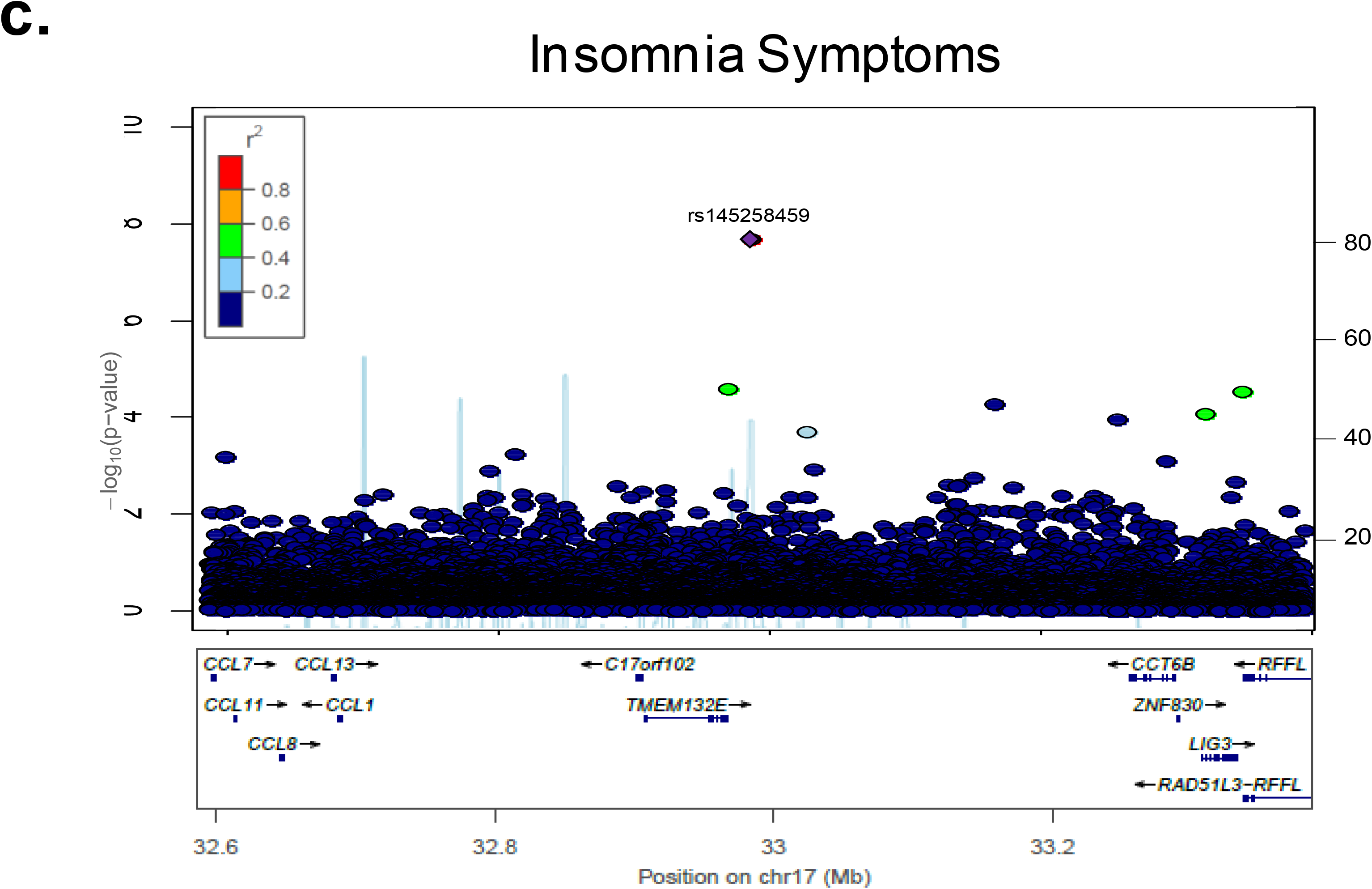
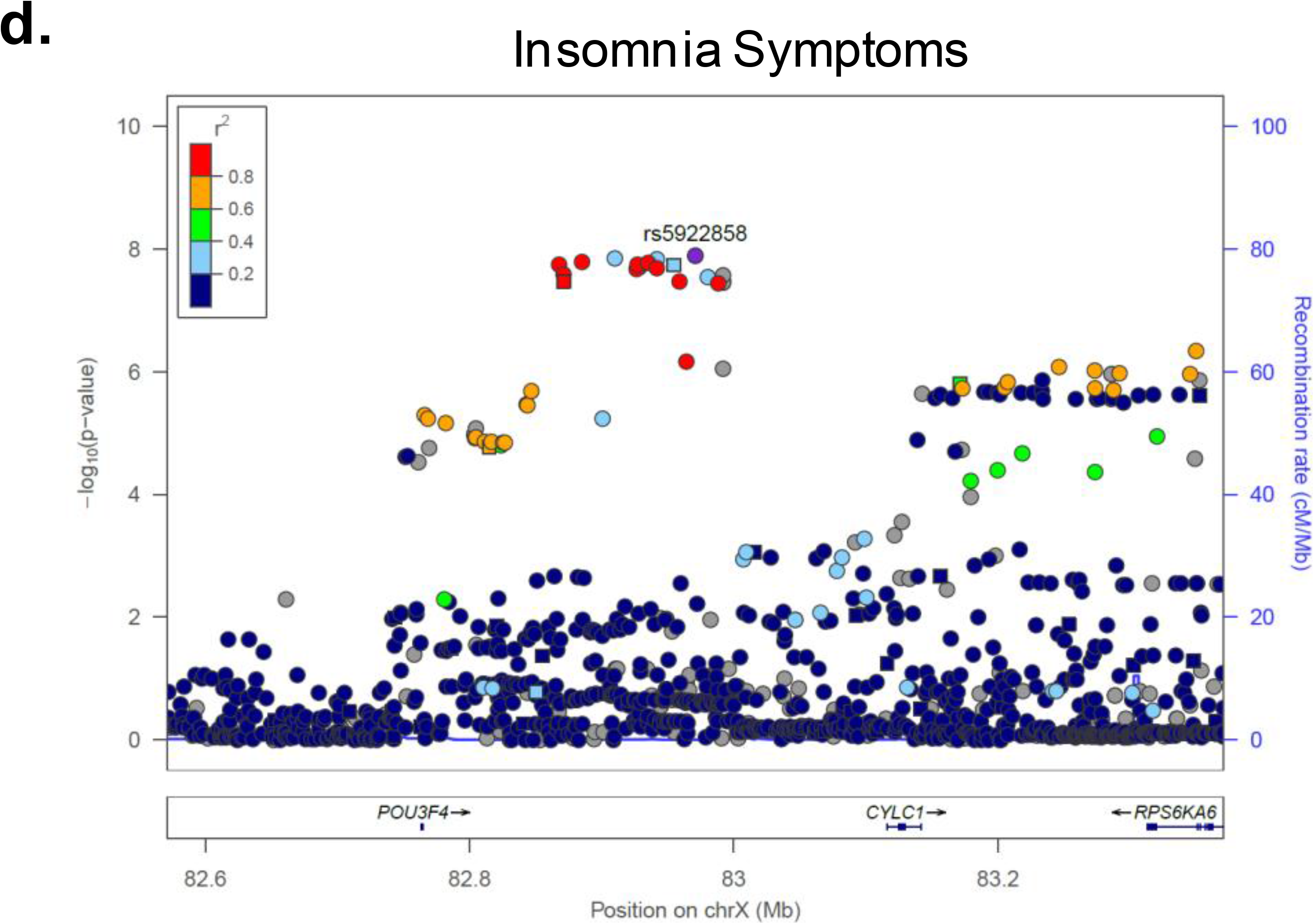
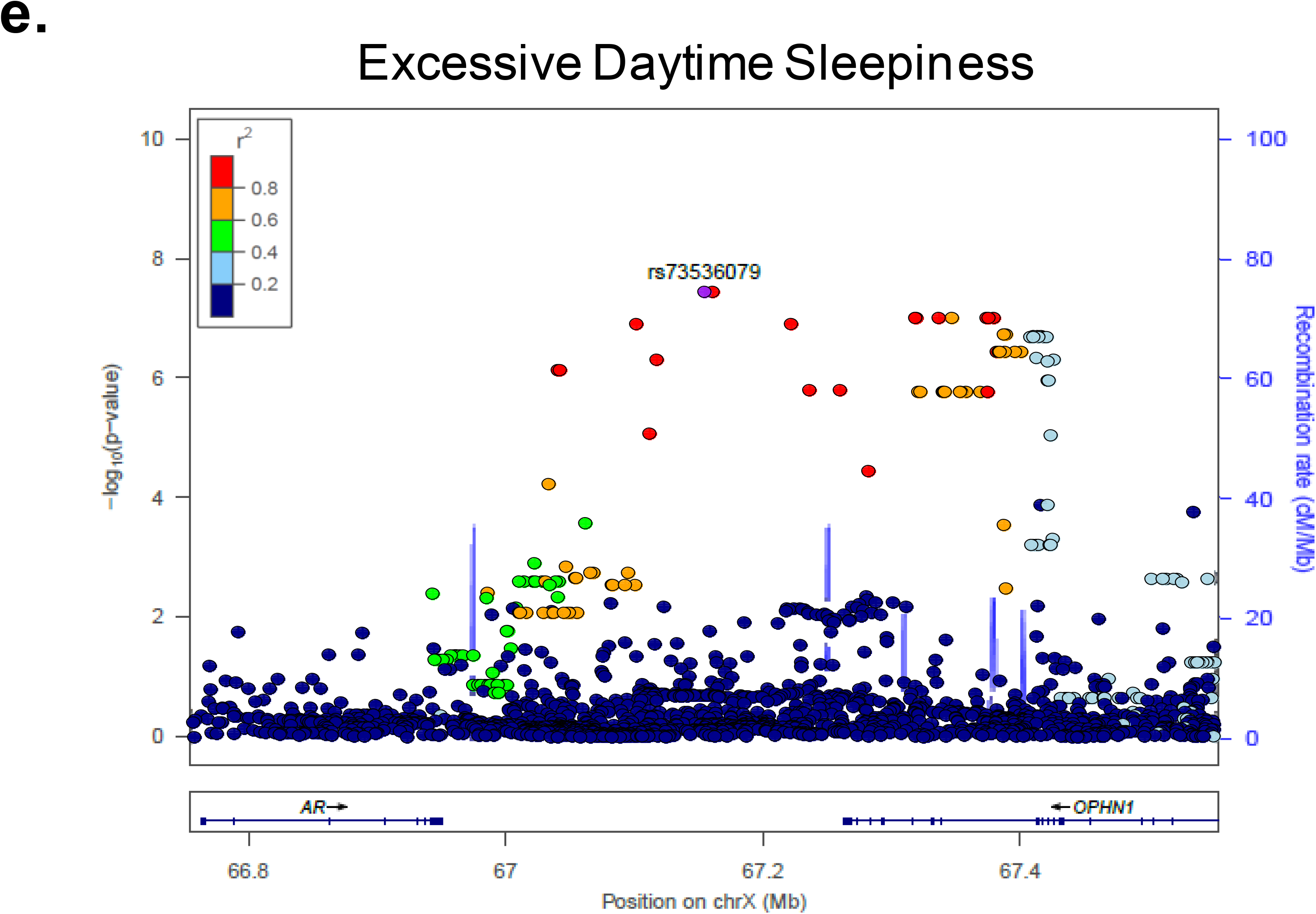
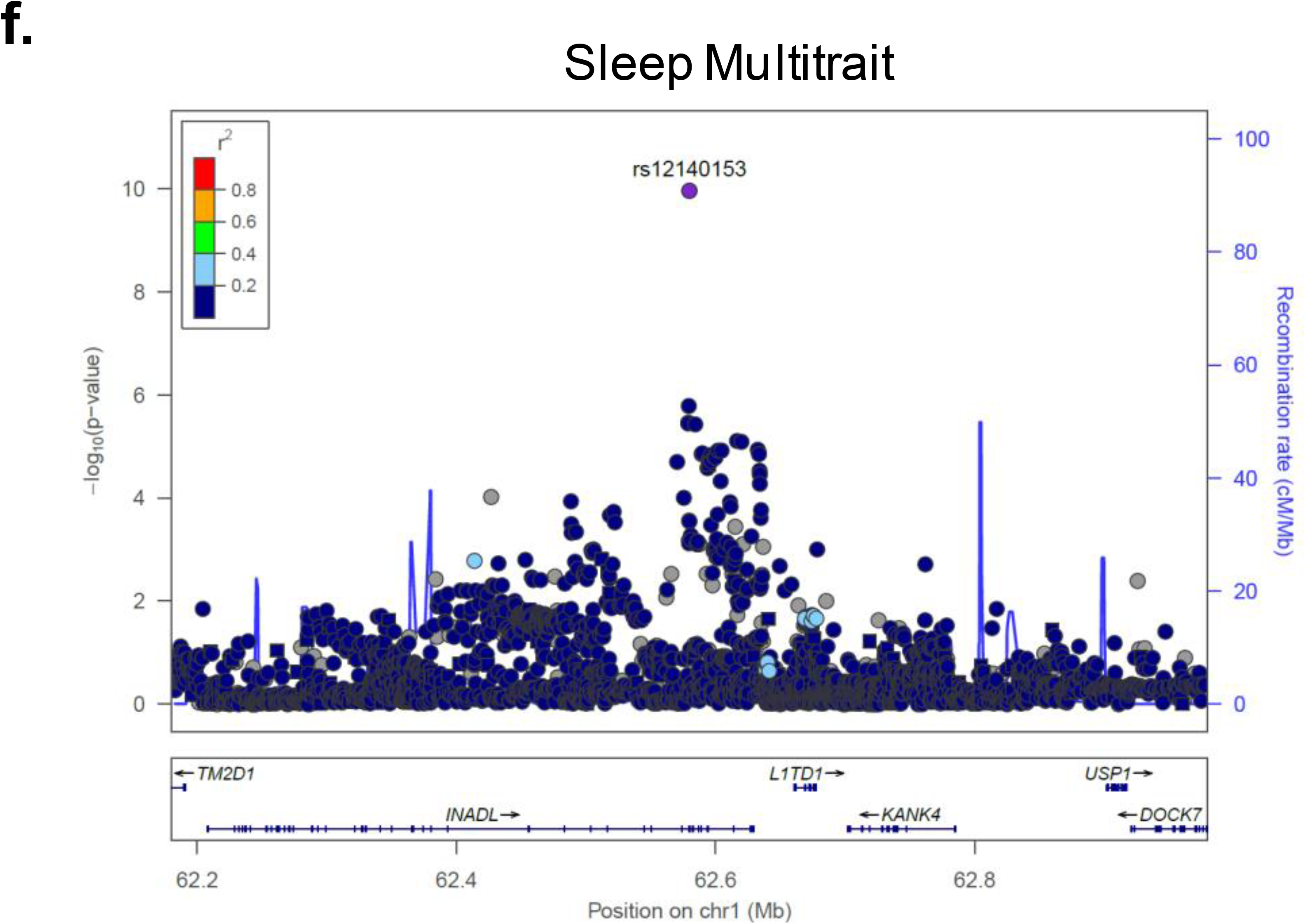
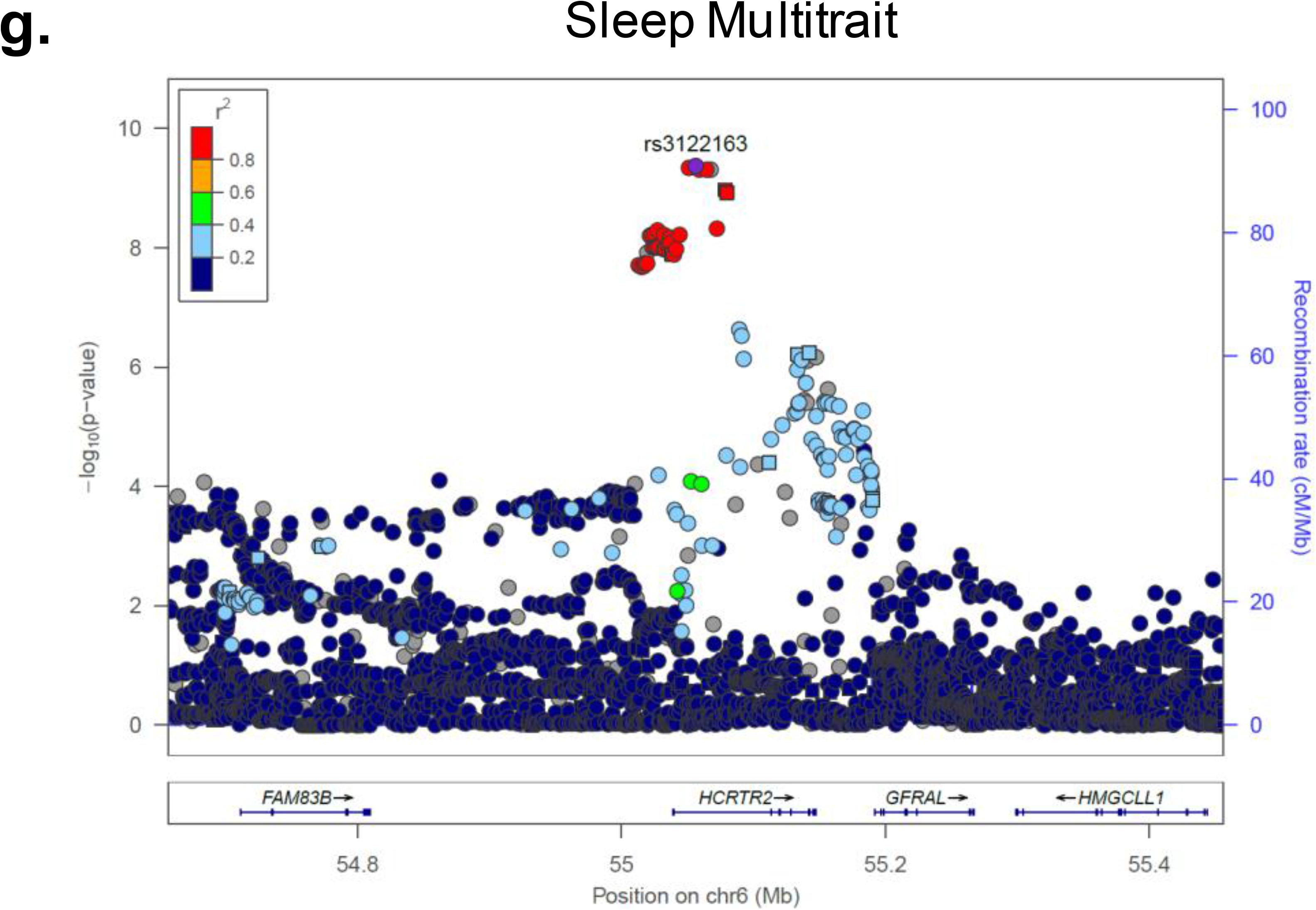
Regional association plots for genome-wide significant loci. Panel **a** sleep duration, **b–d** insomnia symptoms, **e** excessive daytime sleepiness, **f–g** composite trait of sleep duration, insomnia symptoms, excessive daytime sleepiness, and chronotype. Chromosomal position is indicated on the x-axis and -log_10_ *p-*values for each SNP (filled circles/squares) is indicated on the y-axis, with the lead SNP shown in purple (400kb window around lead SNP shown). Genes within the region are shown in the lower panel. The blue line indicates the recombination rate. Additional SNPs in the locus are colored according to linkage disequilibrium (*r*^2^) with the lead SNP (estimated by LocusZoom based on the CEU HapMap haplotypes or within UK Biobank (panel **c**). Squares represent directly genotyped SNPs, and circles represent imputed SNPs.

The leading associations overlap interesting candidate genes enriched in murine/zebrafish hypocretin expressing neurons^37,38^, differentially expressed in sleep-deprived rats^39^, and/or regulate sleep in *Drosophila*^40^. Credible set analyses^41^ highlighted a number of potential causal variants at each locus (**Table 1**) and future experimental studies will be necessary. Bioinformatic annotations^42^ offer an initial opportunity at *in silico* functional interpretation (**Supplementary Table 5; Supplementary Fig. 5**). For example, multiple variants for all three traits are predicted to disrupt binding of *FOXP1*, a neural transcriptional repressor implicated in intellectual disability, autism and language impairment^43^. Interestingly, the *PAX-8* sleep duration association is adjacent to the only chromosomal fusion site since divergence of humans from other hominids ~5 million years ago^44,45^, and the novel genomic structure created by this unique evolutionary history may play a causal role. Pathway analysis^46^ of significant and suggestive loci revealed enrichment of genes associated with immune, neuro-developmental, pituitary and communication disorders (*p*<0.01), and enriched for transcription factor-binding sites for stress-responsive heat-shock-factor 1 (HSF1) and endoplasmic reticulum stress/unfolded protein-responsive factor HERPUD1 **(Supplementary Tables 6&7**).

Aside from the lead *PAX-8* SNPs and a *DRD2* region variant^47^ for sleep duration, limited evidence of association was observed for previously published candidate gene or GWAS signals (*p*_meta_<5x10^−5^; **Supplementary Table 8**), or for regions encompassing core clock genes (**Supplementary Fig. 6**). Our findings for sleep duration GWAS largely overlap with Jones et al.^18^, despite differences in exclusion criteria and analytic approach. Particularly, our study excluded shift workers (n=6,557), sleep medication users (n=1,184) and first-to-third degree relatives (n=7,980), whereas the linear mixed-model analyses by Jones et al. included these populations, leading to a larger sample size (n=127,573). Likely due to this increase in power, Jones et al. identified two additional signals at *VRK2* that did not attain genome-wide significance in our study (rs1380703A β(se)=1.5(0.30) mins/allele and rs17190618T, β(se)=1.60(0.34) mins/allele, *p*=3.8x10^−6^).

Trait heritability calculated as the proportion of trait variance due to additive genetic factors measured here (observed scale SNP heritability, h^2^ (S.E.)) was 10.3 (0.006)% for sleep duration, 20.6 (0.011)% for insomnia symptoms and 8.4 (0.006)% for sleepiness (BOLT-REML variance components analysis^48^). LD-score regression analysis^49^ confirmed no residual population stratification (Intercept (SE): Sleep Duration 1.012 (0.008), Insomnia Symptoms 1.003 (0.008), Excessive Daytime Sleepiness 1.005 (0.007). Tests for enrichment of heritability by functional class using an LD-score regression approach^50^ identified excess heritability across active transcriptional regions for insomnia symptoms and genomic regions conserved in mammals for all three sleep traits. Consistently, heritability enrichment in conserved regions was seen for traits demonstrating significant genetic correlation with sleep (**Fig. 3, Supplementary Table 9**).

**Figure 3.**
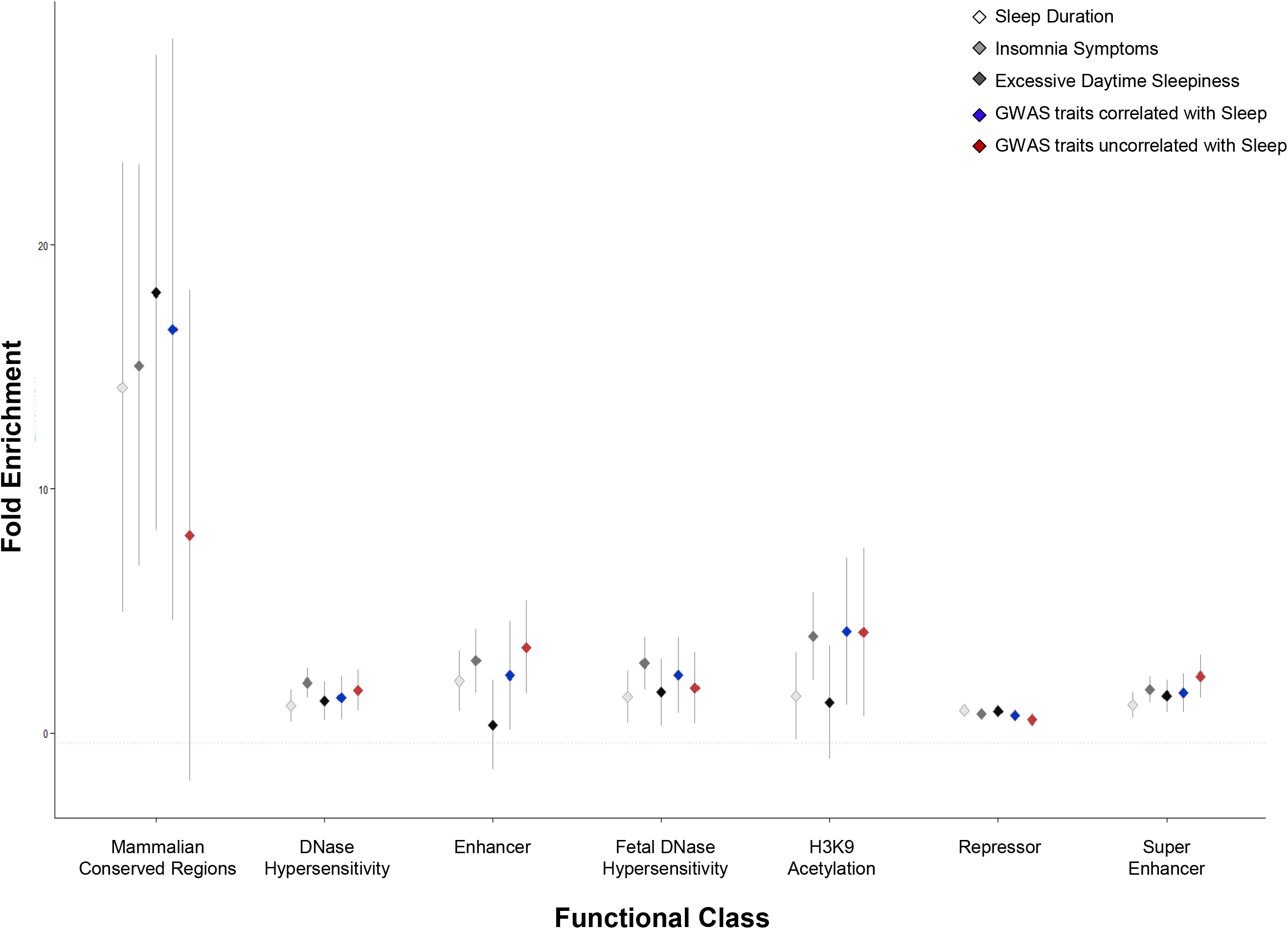
Partitioning of genetic architecture of sleep duration, insomnia symptoms, and excessive daytime sleepiness across functional annotation categories. Fold enrichment estimates for the main annotations of LD-score regression^50^ are indicated on the y-axis across functional annotation class on the x-axis for each trait. Error bars represent the 95% confidence interval around the estimate. 25 functional annotations were tested, and annotations passing the multiple testing threshold (p<0.005) are shown. For context, the average enrichment across functional annotation categories is shown for 9 traits with significant genetic correlation to at least one sleep trait (GWAS traits correlated with Sleep: includes GWAS for BMI, waist circumference, birth weight, depression, educational attainment, three glycemic traits in non-diabetics, and schizophrenia) or for 5 traits with no significant genetic correlation to any sleep traits (GWAS traits uncorrelated with Sleep: includes GWAS for Alzheimer’s Disease, Type 2 Diabetes, autism, rheumatoid arthritis, and height). Abbreviations: H3K9=histone H3 lysine 9.

Sleep duration, insomnia symptoms, excessive daytime sleepiness, and chronotype, are significantly correlated both at the phenotype and genetic level (**Fig. 1**), with greater pair-wise correlations in males as compared to females (Supplementary Fig.1). Thus, in order to find loci common to sleep traits, we performed a multi-trait GWAS^51^. We identified two novel association signals near *HCRTR2* and *INADL*, and revealed that *PAX-8* and *MEIS-1* associations influence multiple sleep traits (**Fig. 2; Table 2, Supplementary Fig. 7**). *HCRTR2* encodes hypocretin receptor 2, the main receptor of two receptors for wake-promoting orexin neuropeptides^52^ involved in narcolepsy and regulation of sleep. Notably, the minor allele at rs3122163 (C) showed sub-threshold association with shorter sleep duration and morningness chronotype, suggesting gain of function, but no association with insomnia symptoms. Assessment of objective sleep measures, functional and physiologic follow-up should yield important insights into orexin receptor signaling, a pathway important for the pharmacological treatment of narcolepsy^53^ and insomnia^54^. *INADL* encodes a membrane protein involved in the formation of tight junctions, and is implicated in photoreception in mice and *Drosophila*^55,56^. The INADL protein is reported to interact with HTR2A^57^, a serotonin receptor with a known role in sleep regulation^58,59^.

**Table 2.**
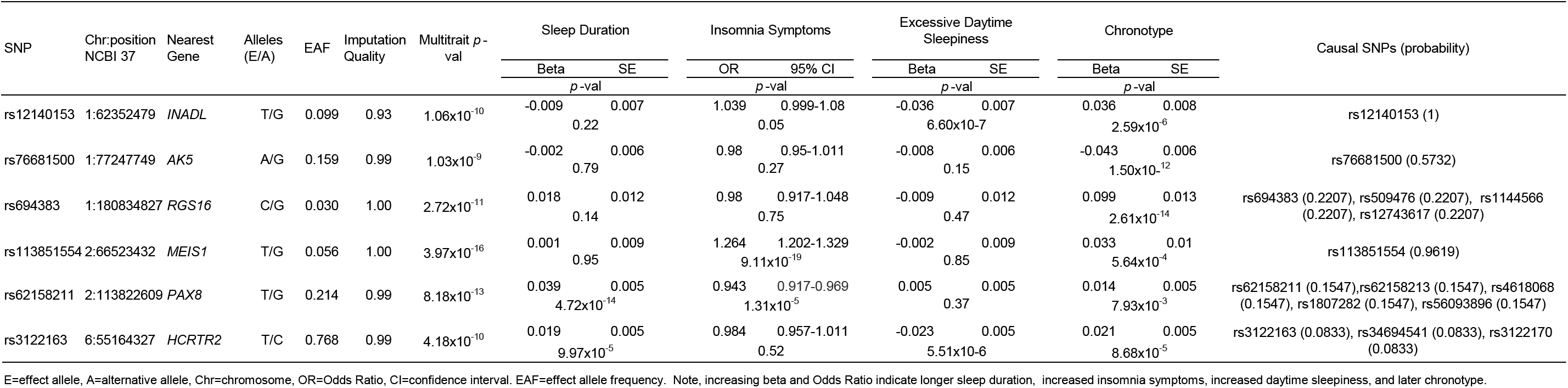
Genome-wide significant (p<5x10^−8^) loci associated with a multiphenotype model of sleep duration, insomnia symptoms, excessive daytime sleepiness and categorical chronotype in subjects of European ancestry in the UKBiobank.

Our strongest association for insomnia symptoms fell within *MEIS1,* a locus previously associated with RLS in GWAS^60^. Our lead SNP rs113851554 and the correlated 3’UTR variant rs11693221 (pair-wise r^2^=0.69, D’=0.90 in 1KG EUR) represent the strongest known genetic risk factor for RLS and were identified in follow-up sequencing studies of MEIS1^61,62^ of the original RLS GWAS signal rs2300478^60,63^. Conditional analysis suggests that only one underlying signal detected by the lead SNP rs113851554 in our GWAS explains the association of all three SNPs with insomnia symptoms (**Supplementary Fig. 8; Supplementary Table 10).** To further investigate the extent of overlap between RLS and insomnia symptoms, we tested if a weighted genetic risk score (GRS) for RLS^64,65^ was also associated with insomnia symptoms with concordant direction of allelic effects (OR [95%CI]= 1.06[1.05-1.07] per RLS risk allele, *p*=1.17x10^−21^; **Supplementary Table 11**). Weighting of RLS GWAS alleles by SNP effects on periodic limb movements (PLMs) did not substantially alter overall results (**Supplementary Table 11**). Interestingly, recent data indicating increased thalamic glutamatergic activity in RLS provides evidence for an underlying propensity for hyperarousal in RLS^66^, which is also a core feature of insomnia. Future analyses of pair-wise bidirectional causal effects for all three traits will be necessary to determine if shared genetic associations represent causality, partial mediation or pleiotropy.

Strong epidemiologic associations of sleep duration, insomnia symptoms and sleepiness have been observed with disease traits, but the extent to which the underlying genetics is shared is unknown. Therefore, we tested for genome-wide genetic correlation between our sleep GWAS and publicly available GWAS for 20 phenotypes spanning a range of cognitive, neuropsychiatric, anthropometric, cardio-metabolic and auto-immune traits using LD-score regression^67^ (**Fig. 4** and **Supplementary Table 12)**.

**Figure 4.**
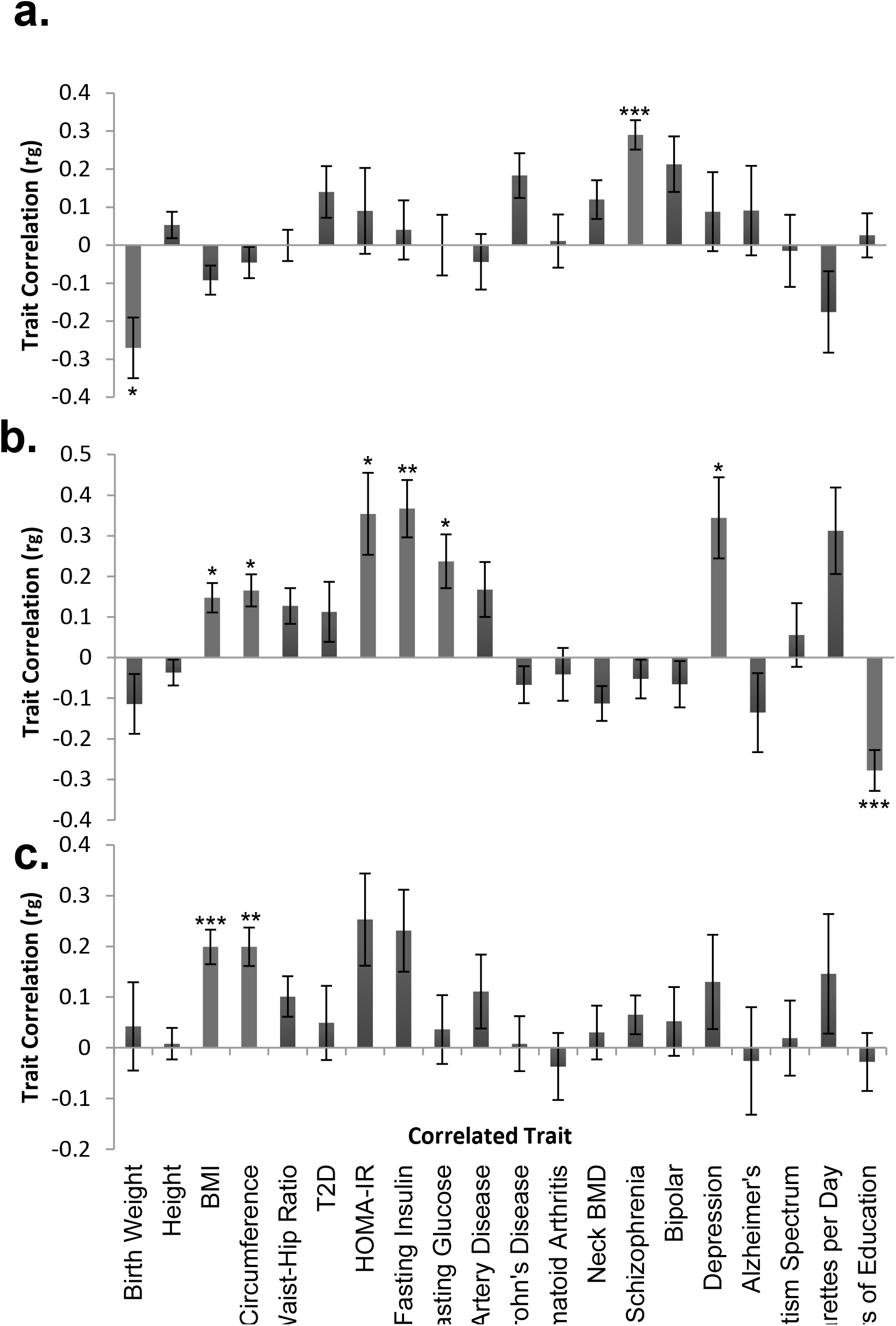
Shared genetic architecture between sleep duration, insomnia symptoms, or excessive daytime sleepiness and 20 behavioral and disease traits. LD-score regression^67^ estimates of genetic correlation (r_G_) of sleep duration (**a**), insomnia symptoms (**b**), and excessive daytime sleepiness (**c**) are compared with the summary statistics from 20 publicly available genome-wide association studies of psychiatric and metabolic disorders, immune diseases, and other traits of natural variation. The x-axis indicates the phenotype compared to each sleep trait and the y-axis indicates genetic correlation (r_G_). Error bars are standard errors. Abbreviations: BMI=body mass index, ADHD=attention deficit hyperactivity disorder, T2D=type 2 diabetes. * p<10^−3^, **p<10^−5^, ***p<10^−7^. After Bonferroni correction, *p*-value cut-off is 0.0025.

Genetic correlations demonstrated a strong biological link between longer sleep duration and risk of schizophrenia (r_G_=0.29, *p*=10^−13^), as suggested by previous reports^18,47,68^. Furthermore, a schizophrenia GRS (96 variants) was associated with longer sleep duration (β(se)=1.44(0.36) mins/allele, *p*=2.56x10^−4^ [2.3 hr inter-quartile range], although a variety of sleep behaviors are seen in schizophrenia patients^69–71^. Significant genetic correlation between low birth weight and longer sleep duration (r_G_= −0.27, p=10^−4^) may reflect shared links between genetically-determined insulin secretion or action pathways underlying fetal growth^72,73^ and long sleep duration. In support, significant genetic correlation was observed by Jones et al.^18^ between over-sleepers and both fasting insulin and risk of type 2 diabetes in UK Biobank. Genetic correlation between sleep duration and Crohn’s disease risk (r_G_=0.18, p=10^−3^) is also consistent with epidemiologic observations^74^.

Significant genetic correlation was also found between increased insomnia symptoms and adverse glycemic traits, increased adiposity and fewer years of education, and between excessive daytime sleepiness and increased adiposity (all p<10^−3^), further highlighting biological overlap of sleep traits with metabolism and educational attainment^17^. In support, studies have shown that experimentally suppressing slow wave sleep leads to decreased insulin sensitivity and impaired glucose tolerance^75,76^. Notably, a fasting insulin GRS was not significantly associated with insomnia symptoms (7 SNPs, OR =1.01, *p*=0.51). Finally, consistent with a well-established but poorly-understood link between excessive daytime sleepiness and obesity^77,78^, a BMI GRS was associated with excessive daytime sleepiness (95 SNPs, β(se) 0.002(0.0004) sleepiness category/allele, *p*=1.67x10^−4^), but not with insomnia symptoms (OR=1.00, *p*=0.73).

Moving forward, replication and systematic testing of genetic correlations in larger samples will be needed. Importantly, genetic correlation testing between insomnia and RLS should be examined, but was not possible here because RLS consortium GWAS results were not available. Additionally, identifying causal relationships between genetically correlated traits may be difficult, and findings using Mendelian randomization approaches will need cautious interpretation given potential selection biases in UK Biobank^79–81^.

In summary, in a GWAS of sleep traits, we identified new genetic loci that point to previously unstudied variants that might modulate the hypocretin/orexin system, and influence retinal or neural development or cerebral cortex genes. Furthermore, genome-wide analysis suggests that sleep traits share underlying genetic pathways with neuropsychiatric and metabolic disease. This work should advance understanding of molecular processes underlying sleep disturbances, and open new avenues of treatment for sleep disorders and related diseases.

## Methods

### Population and study design

Study participants were from the UK Biobank study, described in detail elsewhere^80–82^. In brief, the UK Biobank is a prospective study of >500,000 people living in the United Kingdom. All people in the National Health Service registry who were aged 40-69 and living <25 miles from a study center were invited to participate between 2006-2010. In total 503,325 participants were recruited from over 9.2 million mailed invitations. Self-reported baseline data was collected by questionnaire and anthropometric assessments were performed. For the current analysis, individuals of non-white ethnicity were excluded to avoid confounding effects.

### Sleep quality, quantity and covariate measures

Study subjects self-reported sleep duration, insomnia symptoms, excessive daytime sleepiness, depression, medication use, age, sex, height and weight on a touch-screen questionnaire. For sleep duration, subjects were asked, “About how many hours sleep do you get in every 24 hours? (please include naps)?” with responses in hour increments. To assess insomnia symptoms, subjects were asked, “Do you have trouble falling asleep at night or do you wake up in the middle of the night?" with responses “never/rarely”, “sometimes”, “usually”, “prefer not to answer”. To assess daytime sleepiness, subjects were asked “How likely are you to doze off or fall asleep during the daytime when you don't mean to? (e.g. when working, reading or driving)?" with responses “never/rarely”, “sometimes”, “often”, “all the time”, “don’t know”, “prefer not to answer”. Approximately 500,000 subjects answered these questions, but only the 120,286 unrelated individuals with genetic data and European ancestry were considered for this analysis. Subjects with self-reported shift work (n=6,557) or sleep medication use (n=1,184) were excluded. Subjects who responded “Do not know” or “Prefer not to answer” were set to missing. Sleep duration and excessive daytime sleepiness were untransformed and treated as continuous variables, with daytime sleepiness coded 1–4. The insomnia symptom trait was dichotomized into controls (“never/rarely”) and cases (“usually”). Covariates used in sensitivity analyses included self-reported sleep apnea, BMI, depression, psychiatric medication use, socio-economic, smoking, employment and marital status, and snoring, and secondary GWAS for sleepiness included adjustment for BMI or depression. Sleep apnea cases were defined based on ICD10 diagnosis code (391 cases). BMI at baseline visit was calculated from entries of height and weight (n=75,540 with available data). Depression was reported in answer to the question “How often did you feel down, depressed or hopeless mood in last 2 weeks?” (cases, n=4,242 based on answers “more than half the days”, or “nearly every day”). Medication use was self-reported as part of the initial UK Biobank interview. Our list of psychiatric medication for sensitivity analysis included the four most widely used: fluoxetine (Prozac), citalopram (Cipranol), paroxetine (Seroxat), and sertraline (Lustral). Our list of sleep medications included the 21 most widely used sleep medications in the UK Biobank: oxazepam, meprobamate, medazepam, bromazepam, lorazepam, clobazam, chlormezanone, temazepam, nitrazepam, lormetazepam, diazepam, zopiclone, triclofos, methyprylone, prazepam, triazolam, ketazolam, dichloralphenazone, clomethiazole, zaleplon, butobarbital. Smoking status was self-reported as past smoking behavior and current smoking behavior, and classified into “current”, “past”, or “never” smoked. Socio-economic status was represented by the Townsend deprivation index, based on national census data immediately preceding participation in the UK Biobank. Employment status was self-reported (cases=retired, controls=currently employed). Marital status was derived from self-reported household occupancy and relatedness data. Snoring was reported in answer to the question “Does your partner or a close relative or friend complain about your snoring?”.

### Genotyping, quality control and imputation

Of the ~500,000 subjects with phenotype data in the UK Biobank, ~153,000 are currently genotyped. Genotyping was performed by the UK Biobank, and genotyping, quality control, and imputation procedures are described in detail at the UK Biobank website (http://biobank.ctsu.ox.ac.uk/). In brief, blood, saliva, and urine was collected from participants, and DNA was extracted from the buffy coat samples. Participant DNA was genotyped on two arrays, UK BiLEVE and UKB Axiom with >95% common content. Genotypes were called using Affymetrix Power Tools software. Sample and SNP quality control were performed. Samples were removed for high missingness or heterozygosity (480 samples), short runs of homozygosity (8 samples), related individuals (1,856 samples), and sex mismatches (191 samples). Genotypes for 152,736 samples passed sample QC (~99.9% of total samples). SNPs were excluded if they did not pass QC filters across all 33 genotyping batches. Batch effects were identified through frequency and Hardy-Weinberg equilibrium tests (p-value <10^−12^). Before imputation, 806,466 SNPs pass QC in at least one batch (>99% of the array content). Population structure was captured by principal component analysis on the samples using a subset of high quality (missingness <1.5%), high frequency SNPs (>2.5%) (~100,000 SNPs) and identified the sub-sample of European descent. Imputation of autosomal SNPs was performed to a merged reference panel of the Phase 3 1000 Genome Project and the UK10K using IMPUTE2^83^. Data were prephased using SHAPEIT3^84^. In total, 73,355,677 SNPs, short indels and large structural variants were imputed. X-chromosome data were imputed separately, using Eagle 2.0 for pre-phasing with the -X chromosome flag (no reference panel) in the entire cohort^85^ and IMPUTE2^83^ with the Phase 3 1KG Project reference panel for imputation using the -chrX flag on 500kb chunks in randomly assigned subsets of 30,000 individuals. Post-imputation QC was performed as previously outlined (http://biobank.ctsu.ox.ac.uk/) and an imputation info score cut-off of 0.8 was applied. For GWAS, we further excluded SNPs with MAF <0.001, maximum per SNP missingness of 10%, and maximum per sample missingness of 40%. In total, up to 112,586 samples of European descent with high quality genotyping and complete phenotype/covariate data were used for these analyses.

### Statistical Analysis

Phenotypic correlation analysis was performed using the Spearman test in R using the Hmisc package. Genetic association analysis for autosomes was performed in SNPTEST^86,87^ with the “expected” method using an additive genetic model adjusted for age, sex, 10 PCs and genotyping array. Genome-wide association analysis was performed separately for sleep duration, insomnia symptoms, and excessive daytime sleepiness with a genome-wide significance threshold of 5x10^−8^ for each GWAS. We are 80% powered to detect the following effects: sleep duration β=0.045 hrs (2.7 mins), insomnia symptoms OR=1.07, and excessive daytime sleepiness β=0.021 units (assuming a MAF 0.1, *p=*5x10^−7^) and 80% powered to detect the following effects: sleep duration β= 0.048 hrs (2.9 mins), insomnia symptoms OR=1.08 and excessive daytime sleepiness β=0.023 units (assuming a MAF 0.1, *p=*5x10^−8^). X-chromosome analysis was performed in PLINK 1.9^88^ using linear/logistic regression with separate analysis of the pseudoautosomal regions using the split chromosome flag, adjusting for sex, age, 10 PCs and genotyping array. For the X chromosome signal at rs73536079, we verified using principal components analysis that all carriers of the minor allele fall within the major European ancestry cluster. Follow-up analyses on genome-wide suggestive and significant loci in the primary analyses included covariate sensitivity analysis individually adjusting for sleep apnea, depression, psychiatric medication use, socio-economic, smoking, employment and marital status, and snoring, or BMI (on top of the baseline model adjusting for age, sex, 10 PCs and genotyping array). Sensitivity analysis was conducted only in the subset of subjects with all secondary covariates (n=75,477 for sleep duration, n=39,812 for insomnia symptoms and n=75,640 for excessive daytime sleepiness). Enrichment for disease associated gene sets and transcription factors was performed in WebGestalt^46^ using the human genome as the reference set, the Benjamini Hochberg adjustment for multiple testing, and a minimum number of 2 genes per category. Sex specific GWAS were performed in PLINK 1.9^88^ using linear/logistic regression stratified by sex adjusting for age, 10 principal components of ancestry, and genotyping array. We used a hard-call genotype threshold of 0.1 (calls with greater than 0.1 are treated as missing), SNP imputation quality threshold of 0.80, and a MAF threshold of 0.001. Regional association plots were made using Locuszoom with the HG19 Nov2014 EUR reference panel for background linkage disequilibrium^89^.

Trait heritability was calculated as the proportion of trait variance due to additive genetic factors across the autosomes measured in this study using BOLT-REML^48^, to leverage the power of raw genotype data together with low frequency variants (MAF≥0.001). For multi-trait genome-wide association analysis we applied the CPASSOC package developed by Zhu et al.^51^ to combine association evidence of chronotype, sleep duration, insomnia symptoms and excessive daytime sleepiness. CPASSOC provides two statistics, SHom and SHet. SHom is similar to the fixed effect meta-analysis method^90^ but accounting for the correlation of summary statistics because of the correlated traits. SHom uses a sample size of a trait as a weight instead of variance, so that it is possible to combine traits with different measurement scales. SHet is an extension of SHom but power can be improved when the genetic effect sizes are different for different traits. The distribution of SHet under the null hypothesis was obtained through an estimated beta distribution. To calculate statistics SHom and SHet, a correlation matrix is required to account for the correlation among traits or induced by overlapped or related samples from different cohorts. In this study, we directly provide the correlation matrix calculated from the residuals of four sleep traits after adjusting for age, sex, PCs of ancestry and genotyping array. Post-GWAS genome-wide genetic correlation analysis of LD Score Regression (LDSC)^67^ was conducted using all UK Biobank SNPs also found in HapMap3^89^ and included publicly available data from 20 published genome-wide association studies, with a significance threshold of *p*=0.0026 after Bonferroni correction for all 20 tests performed. As expected, the observed heritability estimates from LDSC^67^ using summary statistics for HapMap3 are lower (5.7 (0.0065)% for sleep duration, 13.3 (0.0123)% for insomnia symptoms and 5.3 (0.005)% for sleepiness) than those calculated by Bolt-REML^48^ using primary data (10.3 (0.006)% for sleep duration, 20.6 (0.011)% for insomnia symptoms and 8.4 (0.006)% for sleepiness), because the HapMap3 panel restricts to variants with >5% MAF. LDSC estimates genetic correlation between two traits from summary statistics (ranging from −1 to 1) using the fact that the GWAS effect-size estimate for each SNP incorporates effects of all SNPs in LD with that SNP, SNPs with high LD have higher X^2^ statistics than SNPs with low LD, and a similar relationship is observed when single study test statistics are replaced with the product of z-scores from two studies of traits with some correlation^67^. Furthermore, genetic correlation is possible between case/control studies and quantitative traits, as well as within these trait types. We performed a weighted genetic risk score analysis using risk scores for restless legs syndrome, schizophrenia, body mass index, and fasting insulin. Risk score SNPs passed the genome-wide significance threshold (p<5x10^−8^) from recent large-scale genome-wide association studies and were present in the UK Biobank (restless legs syndrome 7 SNPs **Supp Table 11**^65^; schizophrenia 96 SNPs^91^; BMI 95 SNPs^92^; fasting insulin 7 SNPs^93^). Independent SNPs were identified and beta estimates recorded for calculation of the weighted risk score. The genetic risk score was calculated by summing the products of the risk allele count multiplied by the effect reported in the discovery GWAS paper. The additive genotype model was used for all SNPs. We performed partitioning of heritability using the 25 pre-computed functional annotations available through LDSC, which were curated from large-scale robust datasets^50^. Enrichment both in the functional regions and in an expanded region (+500bp) around each functional class was calculated in order to prevent the estimates from being biased upward by enrichment in nearby regions. The multiple testing threshold was determined using the conservative Bonferroni correction (p of 0.05/25 classes). Summary GWAS statistics will be made available at the UK Biobank web site (http://biobank.ctsu.ox.ac.uk/).

## Author Contributions

The study was designed by JML, MKR, and RS. JML, JL, IV and RS performed genetic analyses. JML and RS wrote the manuscript and all co-authors helped interpret data, reviewed and edited the manuscript, before approving its submission. RS is the guarantor of this work and, as such, had full access to all the data in the study and takes responsibility for the integrity of the data and the accuracy of the data analysis.

## Acknowledgements

This research has been conducted using the UK Biobank Resource. We would like to thank the participants and researchers from the UK Biobank who contributed or collected data. This work was supported by NIH grants R01DK107859 (RS), R21HL121728 (RS), F32DK102323 (JML), R01HL113338 (JML, SR and RS), R01DK102696 (RS and FS), R01DK105072 (RS and FS), T32HL007567(JL), HG003054 (XZ), The University of Manchester (Research Infrastructure Fund), the Wellcome Trust (salary support for DWR and AL) and UK Medical Research Council MC_UU_12013/5 (DAL). Data on glycemic traits have been contributed by MAGIC investigators and have been downloaded from www.magicinvestigators.org. Data on coronary artery disease / myocardial infarction have been contributed by CARDIo-GRAMplusC4D investigators and have been downloaded from www.CARDIOGRAMPLUSC4D.ORG. We thank the International Genomics of Alzheimer's Project (IGAP) for providing summary results data for these analyses.

The authors have no competing financial interests to declare.

